# Personalized real-time inference of momentary excitability from human EEG

**DOI:** 10.1101/2025.08.31.673404

**Authors:** Lisa Haxel, Oskari Ahola, Jaivardhan Kapoor, Ulf Ziemann, Jakob H. Macke

## Abstract

The efficacy of transcranial magnetic stimulation (TMS) is often limited by non-adaptive protocols that disregard instantaneous brain states, potentially constraining therapeutic outcomes. Current EEG-guided approaches are hindered by their reliance on motor-evoked potentials (MEPs), which confound cortical and spinal excitability and restrict applications to the motor cortex, and a dependence on static biomarkers that cannot adapt to changing neurophysiological patterns. We introduce PRIME (Personalized Real-time Inference of Momentary Excitability), a deep learning framework that predicts cortical excitability, quantified by TMS-evoked potential (TEP) amplitude, from raw EEG signals. By targeting cortical excitability directly, PRIME enables brain state-dependent stimulation across any cortical region. PRIME incorporates transfer learning and continual adaptation to automatically identify personalized biomarkers, allowing stimulation timing to be adapted across individuals and sessions. PRIME successfully predicts cortical excitability with minimal latency, providing a computational foundation for next-generation, personalized closed-loop TMS interventions.

## Introduction

Transcranial magnetic stimulation (TMS) offers a non-invasive approach to modulate cortical excitability and connectivity. Its growing therapeutic applications in neurological and psychiatric disorders include depression, stroke recovery, and movement disorders [1–4]. However, despite widespread clinical adoption, patient outcomes remain highly variable, with more than half of treated individuals showing minimal or no clinical improvement [5–7]. This variability partly stems from a fundamental limitation in conventional TMS design.

Standard protocols operate as “open-loop” systems, delivering predetermined, static patterns of magnetic pulses that remain uniform across sessions and patients. This approach treats the brain as a passive, static recipient of stimulation. Yet, modern neuroscience has established that the brain is better modeled as a dynamic system, with its internal state—characterized by neural oscillations, regional activity, and metabolic demands—varying over coarse-to-fine temporal scales [8]. The neurophysiological and behavioral effects of TMS are therefore governed by principles of state-dependence, where the impact of stimulation depends not only on stimulus parameters, but also on the interaction between the stimulus and the brain’s internal state [9, 10]. Delivering fixed pulse trains into a dynamically fluctuating brain means that therapeutic effects become contingent on the chance alignment of pulses with neurophysiologically receptive moments.

To overcome the limitations of open-loop designs, a system must account for the brain’s excitability state in real time. TMS-evoked potentials (TEPs), recorded via electroencephalography (EEG), directly index local cortical excitability and can be measured from any targeted brain region. Historically, extracting reliable TEPs on a single-trial basis has been challenging due to stimulus-related artifacts and a low signal-to-noise ratio (SNR). Recent advances, however, mitigate these issues by employing individualized, source-level dipole modeling to isolate genuine TEP signals from noise and enable their accurate single-trial extraction [11]. In contrast, motor-evoked potentials (MEPs), recorded peripherally via electromyography (EMG), provide robust measures of corticospinal excitability but are restricted to motor cortex stimulation. The clinical relevance of both biomarkers is underscored by findings that larger TEP and MEP amplitudes predict improved therapeutic outcomes [12–14].

The substantial trial-to-trial variability observed in TEPs and MEPs is driven primarily by fluctuations in pre-stimulus neural activity [15]. First-generation, brain state-dependent TMS protocols aimed to enhance efficacy by timing stimulation to predefined EEG biomarkers, such as particular alpha-band phases or power thresholds [16–18]. Such approaches remain limited by their reliance on fixed biomarkers, which cannot capture the full complexity of moment-to-moment excitability fluctuations. Emerging evidence demonstrates that excitability arises from dynamic, network-level interactions and that its electrophysiological signatures are highly individualized, varying significantly across individuals and sessions [11, 19–23]. While advanced closed-loop protocols propose to overcome the use of fixed biomarkers with adaptive feedback [24], their implementation remains limited by a reliance on MEPs, restricting applications to the motor cortex and precluding their use in clinical contexts that target non-motor regions [25].

To address these gaps—the reliance on fixed biomarkers, restriction to motor cortex applications, and inability to capture individualized excitability patterns—we introduce PRIME (Personalized Real-time Inference of Momentary Excitability), a deep learning framework that predicts TEP amplitudes from raw EEG data at the single-trial level in real-time. Unlike existing systems, PRIME processes EEG signals end-to-end, automatically identifying and adapting to individualized biomarkers that directly reflect cortical responsiveness. Through pretraining on large-scale EEG-TMS datasets and real-time supervised continual learning [**edapt**], PRIME dynamically adjusts to changing brain activity patterns (Fig. 1).

**Figure 1.**
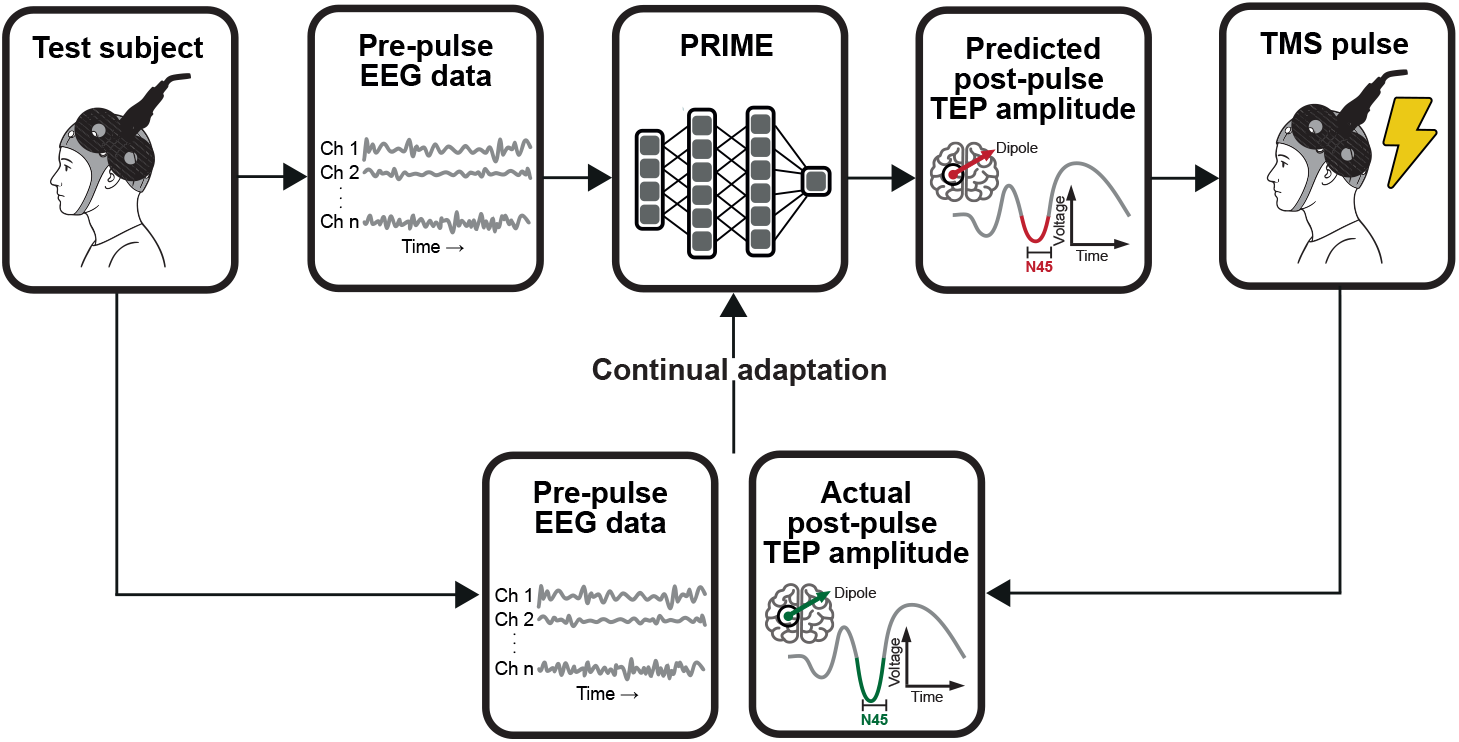
Framework for real-time prediction of TMS-evoked potential amplitudes. Multi-channel electroencephalography (EEG) signals are continuously streamed and preprocessed in real-time. PRIME uses pre-stimulus EEG data to predict the amplitude of the N45 component of the upcoming TMS-evoked potential (TEP) waveform. Immediately following TMS, the actual TEP amplitude is extracted from post-stimulus EEG signals using a current-dipole approach and compared with the prediction. The resulting prediction error, together with recent pre-stimulus EEG data and corresponding TEP markers, are fed back to PRIME for continual updating of the predictive model between pulses.

We demonstrate the real-time predictive performance of PRIME under realistic hardware constraints using previously collected EEG-TMS data. By eliminating dependence on peripheral muscle responses and fixed EEG features, PRIME provides a computational foundation for personalized closed-loop TMS systems applicable across different therapeutic contexts.

## Results

### PRIME accurately predicts cortical excitability in real-time

We developed PRIME to address the limitations of static, open-loop neuromodulation by predicting single-trial cortical excitability directly from pre-stimulus EEG recordings. PRIME uses TMS-evoked potential (TEP) amplitudes as a direct truth measure of cortical responsiveness, providing a computational foundation for personalized closed-loop stimulation. The model’s architecture is designed to process the raw EEG signal from end-to-end, meaning it learns to identify predictive features automatically without depending on predefined biomarkers like alpha- or beta-band power. To achieve this, the model first analyzes local patterns across nearby electrodes and short time windows, then captures how these patterns evolve over longer timescales to learn the complex temporal dependencies within the signal (Fig. 2a).

**Figure 2.**
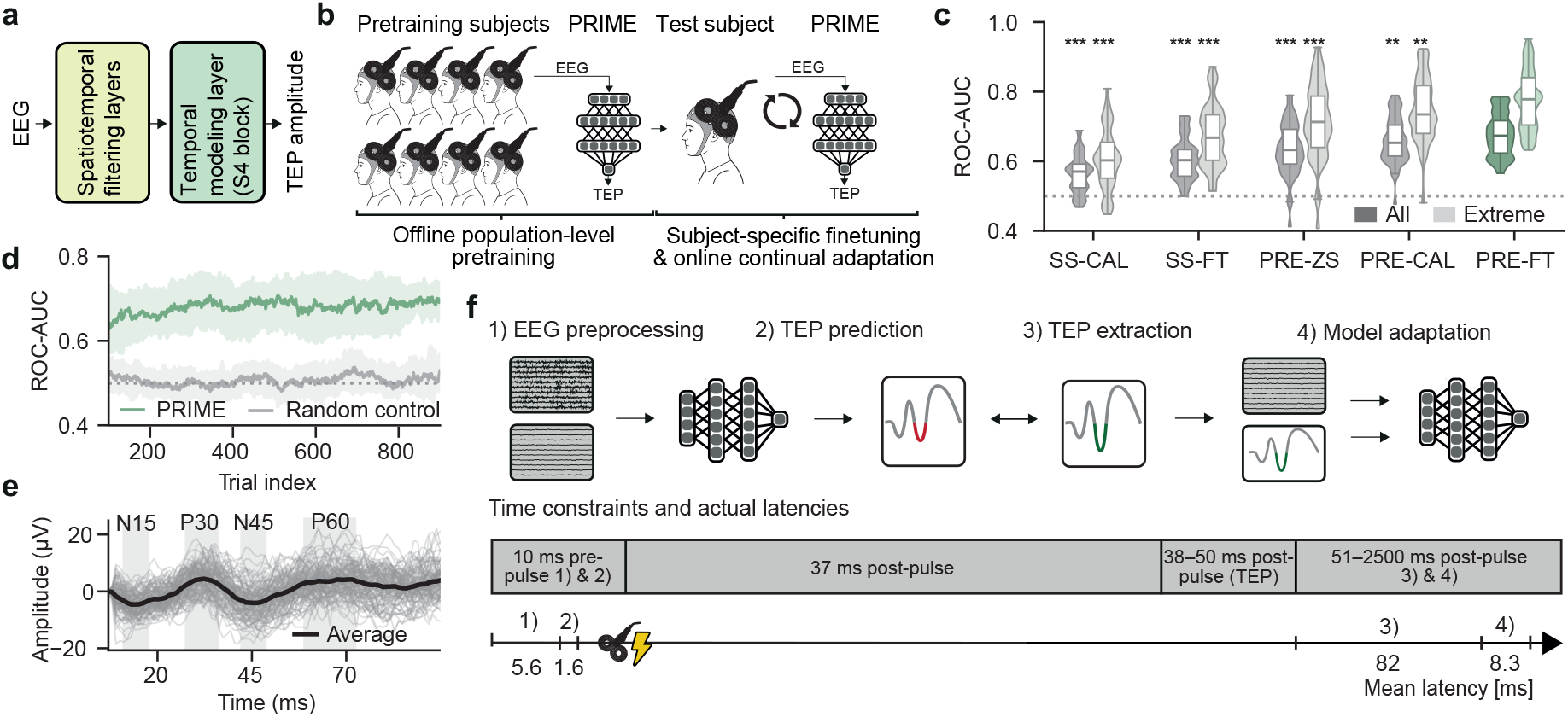
PRIME achieves robust real-time prediction of cortical excitability through multi-stage learning and personalization. **a.** PRIME processes EEG data using spatial-temporal convolutions to identify local patterns in the signal, followed by structured state-space (S4) modeling to capture long-range temporal dependencies and predict upcoming TEP amplitudes. **b.** The training process involves three stages: first, a general model is pretrained on a population of subjects. This model is then personalized for a new subject through an initial calibration, followed by continual real-time adaptation on a trial-by-trial basis. **c.** Predictive performance (ROC-AUC) across 50 datasets (approximately 955 trials each) comparing five training conditions: subject-specific calibration (**SS-CAL**), subject-specific finetuning (**SS-FT**), pretrained zero-shot (**PRE-ZS**, no subject-specific calibration), pretrained calibrated (**PRE-CAL**, calibration without further finetuning), and pretrained finetuned (**PRE-FT**, calibration plus finetuning). Performance is shown for all trials (“All”) and extreme TEP amplitudes (“Extreme”, highest and lowest 25%). PRIME (**PRE-FT**) significantly outperformed all other conditions (** p < 0.01, *** p < 0.001, paired one-sided t-tests, Benjamini-Hochberg corrected). **d.** Predictive accuracy (median ROC-AUC with interquartile range) computed using a sliding window (size = 100) shows rapid model improvement after calibration. PRIME consistently outperformed random controls with shuffled labels. **e.** Representative TEP waveforms from channel F3 (near stimulation site) during the first 100 trials. Individual trials (grey) and their average (black) are shown for the post-stimulus window (10–99 ms). **f.** Computational latency per trial (median across 50 datasets): EEG preprocessing (5.6 ms on CPU), prediction (1.6 ms on GPU), TEP extraction (82 ms on CPU), and model adaptation (8.3 ms on GPU). Pre-pulse processing (7.2 ms) meets real-time constraints (10 ms window), while all post-pulse computations fit within the 2.5 s inter-stimulus interval.

To prepare PRIME for practical use and test its performance, we designed a multi-stage training and adaptation process. Using a 2-fold cross-validation scheme across 50 subjects (954 *±* 149 valid trials per subject), we first trained a general “population model” on data from half the subjects. For each new individual in the other half (the test set), we then calibrated this population model by finetuning it on their first 100 trials, allowing it to adapt to their unique neural characteristics. Finally, during the remainder of the session, the model continually adapted its predictions on a trial-by-trial basis.

We compared the full PRIME system (PRE-FT), which incorporates population-level pretraining, subject-specific calibration, and continual adaptation, against four control conditions to systematically evaluate the contribution of each training stage. To isolate the effects of personalization and ongoing learning, we tested a generic “zero-shot” model using only population-level pretraining (PRE-ZS) and a model that included initial calibration but omitted continual adaptation (PRE-CAL). Finally, to assess the value of population-level pretraining itself, we evaluated two models trained exclusively on single-subject data: one using only the initial calibration trials (SS-CAL) and another that also incorporated continual adaptation (SS-FT).

We assessed performance using ROC-AUC across all trials and, additionally, for trials with extreme TEP amplitudes (highest and lowest 25% of responses) (Fig. 2c, Table 3). The complete PRIME system (PRE-FT) achieved superior predictive accuracy compared to all other conditions, with median ROC-AUC values of 0.68 for all trials and 0.77 for extreme trials. Statistical analysis confirmed that PRIME significantly outperformed all comparison conditions (p < 0.01, paired one-sided t-tests with Benjamini-Hochberg correction). Population-level pretraining provided substantial benefits over training from scratch, with PRE-CAL achieving 0.66 ROC-AUC compared to 0.56 for SS-CAL. Analysis of temporal learning dynamics using a sliding window of 50 trials revealed that PRIME consistently maintained superior performance throughout experimental sessions, substantially outperforming a random control model using shuffled TEP labels (Fig. 2d).

PRIME meets computational latency requirements essential for real-time implementation across all subjects on both GPU and CPU hardware. Latency analysis using an 8-core CPU and NVIDIA 2080Ti GPU revealed mean processing times of 5.6 ms for pre-stimulus EEG preprocessing (CPU; 99th percentile: 7.1 ms) and 1.6 ms for GPU-based TEP prediction (99th percentile: 1.7 ms) or 2.1 ms for CPU-based prediction (99th percentile: 2.4 ms). The combined pre-pulse computation time is 7.2 ms (GPU) or 7.7 ms (CPU)—both, well within the 10 ms constraint required to maintain the reported predictive performance. Post-pulse computations, including TEP extraction (CPU; 82.2 ms, 99th percentile: 101.8 ms) and model adaptation (8.3 ms on GPU, 99th percentile: 9.0 ms; 35.3 ms on CPU, 99th percentile: 40.1 ms), operate comfortably within the 2.5 s inter-stimulus interval typical of single-pulse TMS protocols (Fig. 2e, Fig. S1).

### Structured state-space modeling enables superior temporal feature learning

The PRIME architecture is designed to extract predictive features from multi-channel EEG time series through a combination of spatial and temporal processing components (Fig. 3a). The model begins with temporal convolution to learn time-domain features, followed by spatial convolution across all EEG channels to create spatiotemporal representations. These features are processed through normalization, pooling, and dropout layers before entering the core component: a bidirectional structured state-space (S4) block [26]. The S4 block is optimized for capturing long-range temporal dependencies in neural signals and includes residual connections to facilitate stable training. A linear classifier with global average pooling generates the final TEP amplitude prediction.

**Figure 3.**
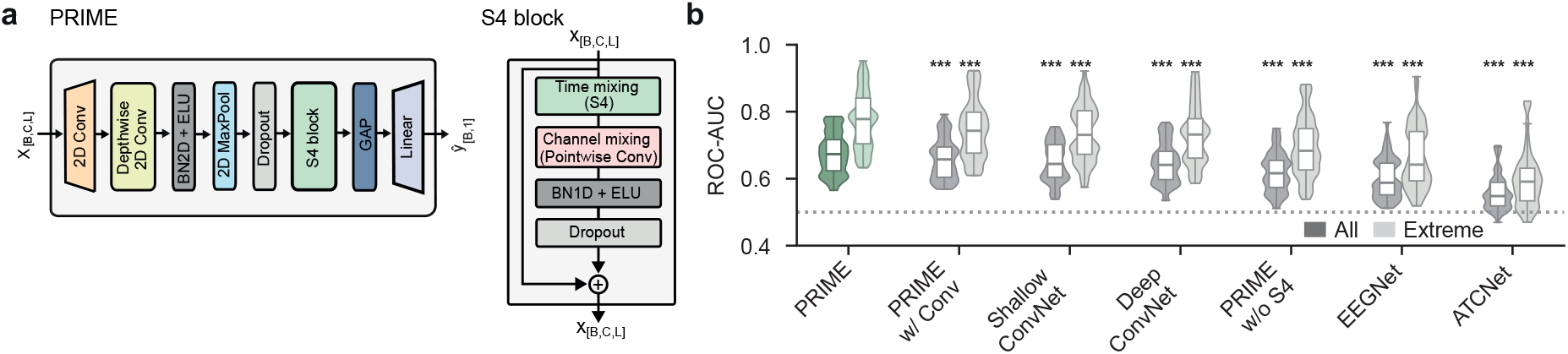
The PRIME structured state-space architecture outperforms conventional EEG decoding models. **a.** PRIME processes input EEG data (*B*, batch; *C*, channels; *L*, time points) through temporal convolution followed by spatial convolution to extract spatiotemporal features. Features pass through normalization, pooling, and dropout layers before entering a structured state-space (S4) block for temporal modeling. A linear classifier with global average pooling generates the final TEP amplitude prediction. **b.** Performance comparison of PRIME against established EEG decoding models (EEGNet, ATCNet, Shallow ConvNet, Deep ConvNet) and ablated variants (PRIME w/o S4; PRIME w/ Conv, i.e., S4 blocks replaced with standard convolution). PRIME significantly outperforms all alternatives for both all trials (“All”) and extreme amplitude trials (“Extreme”, highest and lowest 25%; *** p < 0.001, paired one-sided t-tests, Benjamini-Hochberg corrected). ROC-AUC distributions shown as violin plots across 50 datasets (approximately 955 trials each).

To validate the architectural design of PRIME, we compared performance against four established EEG decoding architectures: ShallowConvNet [27], DeepConvNet [27], EEGNet [28], and ATCNet [29]. We also tested two ablated variants to isolate the contribution of the S4 component: PRIME without the S4 block (PRIME w/o S4), which removes the temporal modeling layer entirely and uses only the convolutional frontend, and PRIME with conventional convolution (PRIME w/ Conv), which replaces the S4 block with standard 1D convolutional layers of equivalent depth.

Across all 50 datasets, the complete PRIME model demonstrated significantly higher predictive accuracy than all comparison models and ablated variants (Fig. 3b, Table 4; p < 0.001, paired one-sided t-tests, Benjamini-Hochberg corrected). The performance degradation in ablated models revealed the critical importance of structured temporal modeling for cortical excitability prediction. Removing the S4 block entirely resulted in a statistically significant performance drop (ROC-AUC 0.62 vs. 0.68 for full PRIME), while replacing S4 with conventional convolution also reduced performance (ROC-AUC 0.66), demonstrating that the superior performance of PRIME stems from the S4 layer’s ability to capture temporal patterns in neural activity that conventional EEG decoding architectures cannot effectively learn.

### Predictive features for cortical excitability are temporally localized

To establish an effective real-time adaptation strategy for PRIME, we compared the performance of several methods (Fig. 4a–d, Tables 5–8). We first established a benchmark using full continual finetuning (CFT), where all model parameters are updated after each trial to allow for full adaptation to the subject’s ongoing brain activity. This method achieved the highest overall predictive accuracy, significantly outperforming all other strategies (p < 0.01 to p < 0.0001).

**Figure 4.**
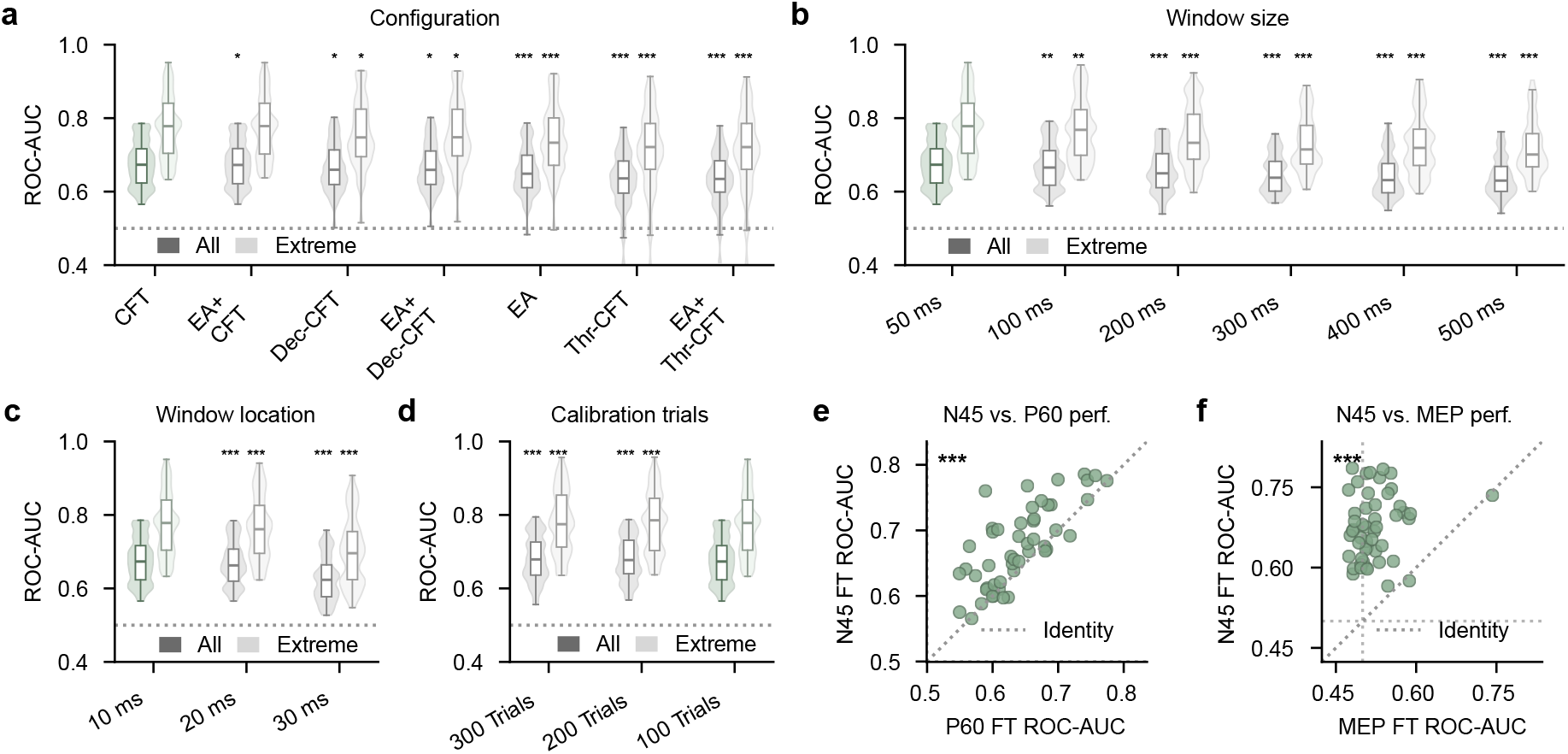
Optimization of PRIME components reveals critical timing constraints and TEP-specific prediction capabilities. a. Performance comparison across PRIME configuration variants: continual finetuning (CFT), Euclidean alignment + CFT (EA+CFT), decision-level continual finetuning (Dec-CFT), Euclidean alignment with decision-level continual finetuning (EA+Dec-CFT), Euclidean alignment only (EA), and thresholded CFT variants (Thr-CFT, EA+Thr-CFT). Standard CFT significantly outperformed most variants for both all trials and extreme trials (highest and lowest 25%; *** p < 0.001, * p < 0.05, paired one-sided t-tests, Benjamini-Hochberg corrected). **b.** Performance is best when using a 50 ms EEG input window. Using longer windows leads to a significant degradation in performance, particularly for extreme trials (** p < 0.01, *** p <0.001, paired one-sided t-tests, Benjamini-Hochberg corrected). **c.** Placing the 50-ms window to end 10 ms before the TMS pulse onset provides significantly better performance than earlier placements (*** p<0.001). **d.** A calibration set of 100 trials per subject is sufficient, larger sets provide only minimal further benefit (** p <0.01, *** p < 0.001, paired one-sided t-tests, Benjamini-Hochberg corrected). **e.** PRIME predicts N45 amplitudes significantly more accurately than P60 amplitudes, with strong cross-subject correlation (*** p < 0.001, paired one-sided t-tests, Benjamini-Hochberg corrected). **f.** PRIME predicts N45 TEP amplitudes significantly more accurately than MEP amplitudes, with no significant correlation between prediction accuracies (*** p < 0.001, paired one-sided t-tests, Benjamini-Hochberg corrected).

We then evaluated whether more targeted, computationally simpler updates could match this performance. One alternative involved updating only the final classifier layer of the model (Dec-CFT), while an even more minimal approach adapted only the decision threshold (Thr-CFT). Both of these strategies resulted in significantly lower accuracy than full CFT, demonstrating that adapting the entire network, including its deep feature-extraction layers, is critical for achieving high performance.

Finally, we tested if regularization could improve performance by preventing the model from drifting too far from its pretrained state. To do this, we incorporated Euclidean Alignment (EA), a technique that aligns the statistical distribution of the incoming data with the population data it was trained on [30]. However, adding EA provided no benefit for predicting extreme trials and significantly lowered performance across all trials. Based on these results, we selected full continual finetuning (CFT) as the most suitable adaptation method for PRIME.

To determine the most effective pre-stimulus EEG window, we investigated the influence of its size and temporal placement on predictive accuracy. We found that performance depends on critical temporal constraints, achieving the highest accuracy with a 50-ms window. Performance declined significantly as we increased the window size from 100 ms to 500 ms (Fig. 4b, Table 6). The placement of this window was equally important; we maximized predictive accuracy by positioning the 50-ms window to end 10 ms before the TMS pulse, as compared to earlier placements (Fig. 4c, Table 7). Finally, when we evaluated calibration requirements, we found that while using 200 or 300 trials provided a statistically significant improvement over a 100-trial baseline, the absolute accuracy gains were minimal (Fig. 4d, Table 8), suggesting that 100 trials provide sufficient subject-specific calibration for practical implementation.

Next, we assessed the ability of PRIME to generalize across different physiological readouts. We found that the model could successfully predict the later P60 TEP component, although its performance was significantly lower than for the N45 component (Fig. 4e, Table 9; paired t-test, p < 0.001). Importantly, we observed a strong correlation between the predictive accuracies for the two TEP components across subjects (Pearson’s r = 0.75, p < 0.0001). In contrast, when we tested whether the model could generalize from cortical to corticospinal excitability (MEPs), we found that its predictive performance for MEPs was substantially lower and approached chance levels (paired t-test, p < 0.001). Consequently, we found no significant correlation between prediction accuracies for the N45 TEP component and MEPs across subjects (Fig. 4f, Table 9; Pearson’s r = 0.17, p = 0.23).

### Theta and alpha oscillations in fronto-central regions drive predictive performance

To identify the neurophysiological features underlying the predictive capabilities of PRIME, we performed occlusion-based sensitivity analysis on subject-specific finetuned models [31]. This approach systematically removes distinct spatial or spectral components from input EEG and measures the resulting decrease in model performance (Δ ROC-AUC), thereby mapping the features most relevant for predicting cortical excitability (Fig. 5). Our analysis revealed that the prediction success of PRIME depends on specific oscillatory patterns localized to discrete cortical regions rather than distributed global brain activity.

**Figure 5.**
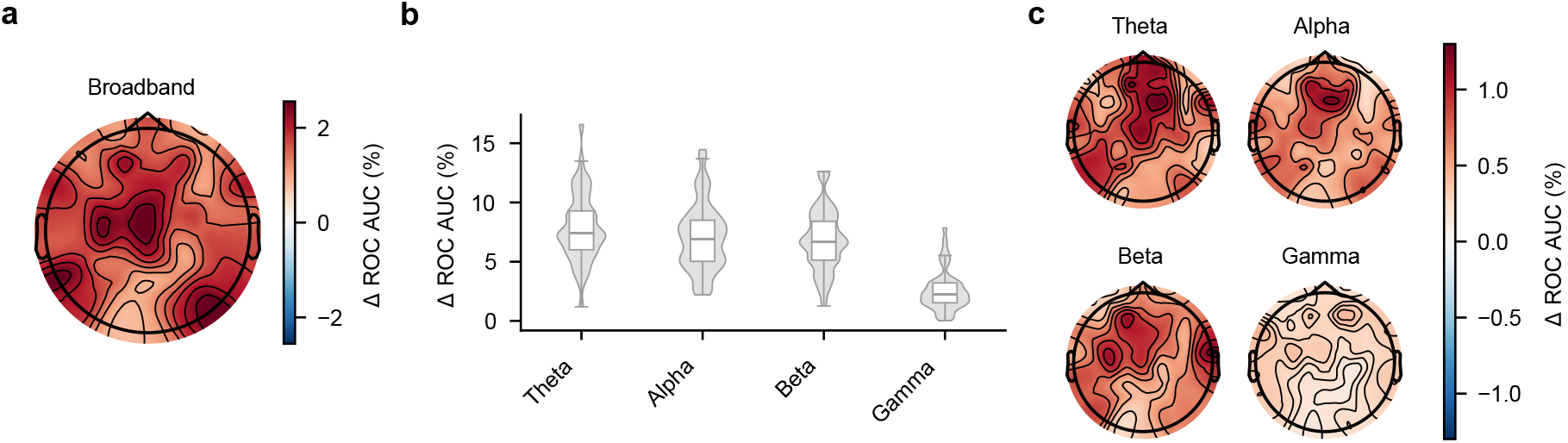
Theta and alpha oscillations in fronto-central regions provide the primary predictive features for cortical excitability. **a.** Spatial distribution of electrode importance determined by occlusion analysis. The magnitude of the decrease in prediction accuracy (positive Δ ROC-AUC values, %) indicates the importance of each electrode and their 8-nearest-neighbors for broadband EEG-based prediction. **b.** Contribution of the theta, alpha, beta, and gamma frequency bands to prediction performance, determined by occluding each band across all electrodes Positive Δ ROC-AUC (%) indicates the importance of each frequency band for accurate prediction. **c.** Spatial distribution of predictive importance for each frequency band, obtained by selectively occluding frequency bands at individual electrodes and their 8-nearest-neighbors. Positive Δ ROC-AUC (%) indicates the importance of specific frequency bands at different cortical locations. All analyses used subject-specific finetuned PRIME models, with statistical significance determined using Wilcoxon signed-rank tests with FDR correction across test subjects (all displayed results significant at p < 0.05).

Spatially, the most predictive broadband EEG signals originated from a localized cluster of electrodes over the left frontal and central cortex (Fig. 5a). Occluding individual electrodes and their nearest neighbors from these regions significantly degraded model performance, whereas electrodes over the right hemisphere contributed minimally to prediction accuracy.

While all frequency bands contained predictive information, the model was most sensitive to low-frequency oscillations (Fig. 5b). Removing theta (4–8 Hz) and alpha (8–13 Hz) bands caused the largest drops in accuracy, followed by beta (13–25 Hz) and gamma (25–47 Hz) bands, respectively.

The predictive power of these oscillations originated from similar cortical areas identified in the broadband analysis (Fig. 5c). The high importance of theta and alpha activity was predominantly localized to left fronto-central regions, with beta band activity showing a similar but weaker spatial pattern. Gamma activity offered negligible predictive value from any specific region.

## Discussion

We introduced PRIME, a deep learning framework designed to predict cortical excitability from real-time EEG data using TMS-evoked potential (TEP) amplitudes as a measure of cortical excitability. Unlike current approaches that rely on static biomarkers, PRIME adapts in real-time to dynamic and individual-specific neural states. This approach addresses critical shortcomings of current EEG-guided TMS: it automatically learns personalized biomarkers instead of using fixed features, continuously adapts to evolving brain states, and targets cortical rather than corticospinal excitability. This TEP-based approach enables closed-loop stimulation across diverse cortical targets for neurological and psychiatric disorders.

PRIME achieved strong predictive performance with a median ROC-AUC of 0.68 across all TEP amplitudes and 0.77 when distinguishing extreme excitability states (highest and lowest 25% of trials). This performance was achieved through the general-to-individual training strategy of PRIME, involving population-level pretraining, subject-specific calibration, and continual online adaptation—which was essential for navigating significant inter-subject variability [6, 7, 23]. By integrating structured state-space (S4) modeling, PRIME captured EEG dynamics relevant to cortical responsiveness, proving superior to established EEG-based decoding networks (Fig.2b, Table4). [32, 33].

PRIME predicts the brain’s immediate electrophysiological response to a single pulse of TMS as an instantaneous read-out of cortical excitability. Instantaneous response amplitude has long served as a surrogate for membrane excitability [6, 34]. Crucially, a more excitable, or receptive, state is not only expected to produce a stronger immediate response but is also considered important for inducing therapeutic long-term changes in synaptic strength (i.e., plasticity) [6, 35]. Repetitive stimulation (rTMS) is thought to induce these plastic changes through molecular cascades similar to long-term potentiation (LTP) [2]. Cortical excitability itself fluctuates rhythmically, most prominently in theta (4–8 Hz) and alpha (8–13 Hz) bands. These rhythms reflect cyclical changes in neuronal excitability. During a high-excitability phase, neurons are closer to their firing threshold and thus more susceptible to external stimulation, creating conditions favorable for inducing LTP [36]. Indeed, delivering TMS at EEG-derived high-excitability phases in humans enhances both the immediate response amplitude and the longer-term, LTP-like after-effects of stimulation [16–18].

PRIME leverages this state-dependence by predicting effective stimulation moments from real-time EEG data. Rather than delivering stimulation randomly with respect to brain state, PRIME ensures that more pulses arrive when neurons are neurophysiologically primed for plasticity induction. This transforms TMS from a probabilistic intervention into a more deterministic therapeutic tool. Potential implications become evident when considering practical scenarios. In a typical 800-trial session, PRIME correctly predicts cortical excitability in 544 trials overall (68% accuracy) and 308 out of 400 extreme excitability trials (77% accuracy), compared to 400 and 200 expected by chance, respectively. This represents a 36% improvement in overall accuracy (54% for extreme states), potentially translating to more efficient stimulation protocols and enhanced therapeutic outcomes.

The neurophysiological signatures underlying the predictive success of PRIME provide further insight into potential cortical excitability mechanisms. Theta (4–8 Hz) and alpha (8–13 Hz) oscillations emerged as the most predictive features (Fig. 4b–c), consistent with their established roles in motor cortical excitability and sensorimotor integration: Theta oscillations coordinate excitatory-inhibitory cycles that support selective information exchange through phase-coupling [37], as demonstrated in both cognitive [38–40] and sensorimotor tasks [41, 42]. Alpha oscillations are associated with voluntary motor control, including imagery, preparation, execution, and somatosensory performance [43–48]. Predictive information was primarily concentrated in fronto-central regions (channels Cz, CPz, CP3, C3, F1), aligning with previous research [11] and reflecting integrative mechanisms linking sensorimotor coordination with cognitive states including attention, alertness, and emotional processes [49–52]. The temporal specificity of predictive features—with optimal performance achieved using a 50-ms window ending 10 ms before stimulation (Fig.3bc, Table6,7)—indicates that cortical excitability states are determined by very recent neural activity patterns. This finding has practical implications for closed-loop implementation, suggesting that real-time systems can rely on relatively short EEG epochs for accurate predictions.

A critical finding of our study is the stark contrast between the ability of PRIME to predict TEP amplitudes versus MEP responses. While achieving robust performance for TEP prediction, the model’s accuracy for MEPs approached chance levels (0.52 ROC-AUC), with no significant correlation between TEP and MEP prediction performance across subjects. This difference suggests fundamental differences between cortical and corticospinal excitability mechanisms. MEP amplitudes likely reflect the integrated output of multiple neural processing stages: cortical excitability, corticospinal tract integrity, spinal motor neuron pool excitability, and neuromuscular transmission [53, 54]. In contrast, TEPs provide a direct measure of cortical responsiveness by capturing post-synaptic potentials evoked by TMS without spinal confounds [55]. The success of PRIME in predicting both N45 and P60 TEP components, with similar spatial and frequency patterns of predictive features (Fig. S2), would be consistent with cortical excitability mechanisms underlying both responses. The slightly lower prediction performance for P60 compared to N45 (0.64 vs. 0.68 ROC-AUC for all trials; 0.72 vs. 0.77 for extreme trials) potentially reflects increased contamination from sensory activity at longer latencies [56] rather than fundamentally different predictive mechanisms. This finding suggests that PRIME is successfully isolating and decoding specific cortical dynamics that are upstream of, and distinct from, factors that modulate spinal and muscular excitability. The clear dissociation indicates that the EEG features PRIME uses capture instantaneous cortical states rather than non-specific markers of global arousal or muscle artifacts.

From a practical implementation perspective, PRIME meets stringent latency requirements for real-time TMS applications on both GPU and CPU hardware, making the approach feasible for standard clinical computing environments. The framework processes EEG data in 7.2 ms on GPU or 7.7 ms on CPU—both well below the 10 ms constraint required to maintain predictive performance without interfering with stimulation timing. While GPU hardware accelerates model updates (8.3 ms vs 35.3 ms), both platforms complete adaptation following TEP extraction (82.2 ms) well within the typical 2.5-second inter-stimulus interval, enabling continuous model refinement without disrupting established TMS protocols.

Several challenges must be addressed before clinical deployment: First, the current evaluation used datasets from controlled laboratory settings. The performance of PRIME must be validated in real-world clinical environments where EEG data are often contaminated by movement artifacts, muscle activity (EMG), and fluctuations in patient vigilance and alertness [57], and the system needs to be investigated in such higher-noise environments. Second, the model was trained on a cohort of healthy young adults, and its ability to generalize across heterogeneous patient populations varying in age, medication status, and comorbidity needs to be established. The neurophysiological signatures of excitability may differ significantly in older adults or patients with different psychiatric comorbidities. Third, while grounded inevidence [16–18], the assumption that consistently stimulating during high-or low-excitability states leads to greater therapeutic LTP or LTD requires direct clinical confirmation. The relationship between excitability and plasticity is likely non-linear, with a potential optimal window rather than a simple “more is better” function [9]. Future clinical trials are needed to measure both symptom improvement and neurophysiological data confirming the engagement and enhancement of LTP-like plasticity.

PRIME establishes a computational framework for personalized neuromodulation by shifting from static biomarkers to dynamically inferred EEG-derived brain states. Its ability to predict cortical excitability with 36-54% improvement over chance levels provides a foundation for more precise, individualized closed-loop TMS interventions. Future clinical applications could leverage the continuous, graded predictions of PRIME to reduce treatment session requirements and enhance therapeutic effectiveness across neurological and psychiatric disorders. The next critical steps involve validating these promising results to translate this technical proof-of-concept into measurable clinical benefits. The real-time adaptability of our framework opens possibilities for advanced closed-loop systems that could dynamically modulate both stimulation timing and intensity to achieve optimal biological effects across fluctuating brain states.

## Acknowledgments

This work was supported by the Else Kröner Medical Scientists Kolleg Clinbrain: Artificial Intelligence for Clinical Brain Research, the Machine Learning Cluster of Excellence, EXC number 2064/1–390727645, the German Federal Ministry of Education and Research (BMBF): Tübingen AI Center, FKZ: 01IS18039A and the European Research Council (ERC Synergy) under the European Union’s Horizon 2020 research and innovation program (ConnectToBrain, grant number 810377)

## Author contributions

Conceptualization: L.H., U.Z., J.H.M., Methodology: L.H., O.A., J.K. Software: L.H., O.A. Validation: L.H., Formal analysis: L.H., Data curation: O.A., Visualization: L.H., Writing – Original draft: L.H., Writing – Review and editing: J.K., O.A., O.P.K., U.Z. and J.H.M., Supervision: U.Z., J.H.M., Project administration: U.Z., J.H.M., Funding acquisition: U.Z., J.H.M

## Methods

### Participants and data acquisition

We analyzed single-pulse TMS–EEG data acquired as part of the European Research Council-funded ConnectToBrain project (grant no. 810377). The data were collected at two research centers (University of Tübingen, Germany and Aalto University, Finland) and comprise recordings from 50 healthy, right-handed adults (26 female; mean age 27 ± 6 years).

The original study protocol was approved by the ethics committees of the University of Tübingen (810/2021BO2) and the Helsinki University Hospital (HUS/1198/2016) and was conducted in compliance with the Declaration of Helsinki. All participants provided written informed consent.

The dataset contains concurrent TMS–EEG and electromyography (EMG) recordings. The experimental protocol consisted of four blocks of 300 biphasic single pulses (1200 total) targeted at the motor hotspot of the left primary motor cortex (M1). Stimulation was delivered at an intensity of 110% of the resting motor threshold (rMT), with a randomized inter-trial interval of 4.0–4.5 s (Aalto) or 2.5–3.5 s (Tübingen). High-density EEG data were recorded with TMS-compatible NeurOne systems (64-channel at Aalto, 128-channel at Tübingen; Bittium Ltd, Finland) at a sampling rate of 5 kHz. EMG was recorded from the right abductor pollicis brevis (APB) and first dorsal interosseus (FDI) muscles.

Site-specific equipment included a Nexstim NBS 5.2.4 or NBT 2.2.4 system (Nexstim Plc, Finland) with a Nexstim cooled coil at Aalto, and a MagVenture R30 or X100 stimulator (MagVenture Inc, United States) with a Cool-B65 coil at Tübingen. Neuronavigation was performed using individual T1-weighted magnetic resonance images (MRIs) with Nexstim (Aalto) or Localite (Tübingen; Localite GmbH, Germany) systems. To mask the TMS click sound, individually calibrated noise was delivered via ER-3C insert earphones (Etymotic Research Ltd, United States). T1-weighted MRIs were acquired using MPRAGE sequences with 3T Siemens Skyra or Prisma scanners.

### Data preprocessing

Our preprocessing pipeline was designed to simulate a real-time analysis workflow and consisted of two primary stages: an offline *calibration* stage to define subject-specific parameters and a simulated online *application* stage where these fixed parameters were applied to subsequent trials.

#### Calibration phase

The first 125 trials per participant were initially selected to ensure a minimum of 100 valid trials post-rejection, from which all preprocessing parameters were derived. The trial count was increased until at least 100 valid trials were acquired.

##### Downsampling and epoching

EEG data, originally sampled at 5 kHz, were polyphase downsampled to 1 kHz for pre-stimulus ([−505, −5] ms) and post-stimulus ([−30, 100] ms) epochs separately. EMG data ([−500, 200] ms) were likewise downsampled to 1 kHz.

##### Channel rejection

Bad channels were iteratively rejected from both pre-and post-stimulus data independently. Pre-stimulus criteria included z-scored median absolute deviation (thresholds [−3, 3]), high-frequency power (30–47 Hz; threshold *>* 5), and lag-1 autocorrelation (thresholds [−4, 4]). Post-stimulus channels were rejected based on the area under the curve in the average response (20–60 ms; threshold *>* 3). Rejected channels were reconstructed using spherical spline interpolation.

##### Independent component analysis (ICA)

Ocular and muscle artifacts were identified using Infomax ICA [58] on pre-stimulus data ([−1100, −5] ms) that was bandpass-filtered (1–100 Hz, 2nd-order Butterworth). ICA was done using the minimum number of principal components explaining *>* 99% of the variance. Ocular components with a probability *>* 0.9 (via ICLabel [59]) were removed; this threshold was lowered to 0.7 if fewer than two components were initially identified (often two components exist; one for blinks and one for lateral eye movement). Trials with high ocular activity (z-score *>* 2 in the IC time series) in the post-stimulus interval ([15, 100] ms) were rejected.

##### Pre-stimulus EEG preprocessing

Data were bandpass-filtered (2–47 Hz, 2nd-order Butterworth), average-referenced, and subjected to iterative trial rejection based on median absolute deviation (z-score thresholds [−8, 4]) and single-channel deviation (z-score threshold */pm*5).

##### Post-stimulus EEG preprocessing

Prior to ICA, the data were baseline-corrected ([−25, −15) ms), the pulse artifact window ([−14, 14) ms) was tapered using a 1-hanning window, and average referenced after interpolating bad channels. Following ICA, data were again baseline-corrected ([−25, −15) ms), average-referenced, and cropped to *t >* 0 ms. We suppressed non-neural artifacts using the source-utilizing noise-discarding (SOUND) algorithm [60] (*λ* = 0.01, tolerance= 10^−9^) and TMS-related muscle artifacts with the fixed-component Signal-Space Projection and Source-Informed Reconstruction (SSP–SIR) algorithm [61]. For SSP-SIR, we removed components explaining 99% of the variance in supra-100-Hz-filtered average data weighted by the artifact kernel estimated using 50-ms sliding windows. Then, an extended TMS pulse artifact window ((0, 15) ms) was tapered. Finally, trials were iteratively rejected based on global (median absolute deviation z-score *>* 5) and local (single-channel z-score *±*5) amplitude criteria in the [20, 60] ms window.

##### EMG preprocessing

EMG data were high-pass filtered (2 Hz, 4th-order Butterworth). Powerline noise was removed by iteratively subtracting 50 and 100-Hz sine fits calibrated on the pre-stimulus interval ([−200, −15] ms). Trials with pre-stimulus muscle activation (min-max difference *>* 50 *µ*V) were rejected. For analysis, we selected the single EMG channel with the highest average MEP amplitude ([20, 50] ms).

#### Application phase and real-time compatibility

The application phase was designed to emulate online, trial-by-trial processing. All preprocessing parameters, including channel interpolation matrices, trial rejection statistics, and frequency-based, ICA, SOUND, and SSP–SIR filters, were fixed during the calibration phase and applied without modification to all subsequent data. This approach ensures that TEP amplitude estimates are not biased by changing preprocessing characteristics over time. The computational latency of each step in this pipeline was assessed to confirm its compatibility with real-time experimental requirements.

### TEP component extraction

To analyze transcranial magnetic stimulation (TMS)-evoked potentials (TEPs), we constructed individualized spatiotem-poral filters following [11] based on current dipole modeling [62, 63]. This approach mitigates the low signal-to-noise ratio (SNR) and various neural and non-neural artifacts present in single-trial TEPs. Our analysis focused on the N45 component, thought to reflect the balance of cortical excitation and inhibition [55]. We defined a target time interval of **T** = [38, 50] ms post-TMS to avoid contamination of early-latency TMS-related artifacts and later-latency sensory-evoked activity [11, 56].

#### Dipole estimation

Individualized dipole models were derived from the first 100 valid calibration trials. For each time point *t* ∈ **T** in the averaged EEG data 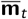, we estimated the optimal dipole position 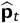 on the cortex and its corresponding moment 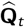 as

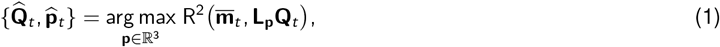

where **L_p_** is the leadfield matrix for a dipole at position **p**, and 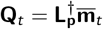 is the single-dipole solution obtained via the Moore-Penrose pseudoinverse. The leadfield was derived from the fsaverage brain template [64], comprising 20,484 sources, and mapped to EEG sensors using the standard_1005 electrode locations.

#### Individualized time-window selection

To account for inter-individual variability in response timing, we defined a personalized, stable, high-amplitude time window 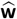 for each participant. This window was identified by penalizing deviations in dipole position and orientation while rewarding high signal amplitude as

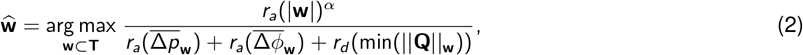

where the window duration *|***w***|* was constrained to be between 3 and 6 ms. In the equation, *||***Q***||***_w_** represents the dipole amplitudes, while 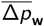 and 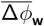 denote the average positional and angular dipole deviances from the mean within window **w**. The function *r_a/d_* (·) ranks windows in ascending or descending order, and *α* was set to 1.5. The final dipole filter position 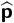 and orientation 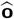 were defined as the amplitude-weighted average over the optimal window 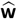.

#### Single-trial amplitude extraction

For each trial *i*, we estimated the dipole amplitude 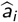 using two approaches. For the **fixed-orientation** model, the scalar amplitude was estimated as

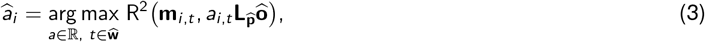

where *a*_*i*,*t*_ is the scalar solution to the least-squares problem 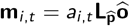. Given that the estimated scalar amplitude can be negative due to noise or polarity reversal, we used the absolute amplitude as the final measure of cortical excitability. For the **free-orientation** model, the amplitude was estimated as

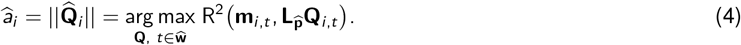

A similar procedure was performed for the P60 component within **T** = [55, 70] ms, using a window duration constraint of 4 ≤|**w***|* ≤ 7 ms.

### Target label generation

We implemented a causal, adaptive normalization procedure to transform single-trial TEP and MEP amplitudes into a stable excitability metric that is robust to slow temporal drifts. This process mirrored the main pipeline, with a calibration phase to derive statistics and an application phase to generate labels.

#### Calibration: Normalization parameter estimation

During the calibration phase, we derived a fixed set of normalization parameters using the single-trial amplitudes extracted from the same set of initial trials used for the main preprocessing (≥ 100 valid trials). First, this amplitude time series was causally detrended using an Exponentially Weighted Moving Average (EWMA) with a 25-trial span. To ensure stable parameter estimation, the initial 25 trials were disregarded to account for the filter’s warm-up period. From the remaining stable, detrended values, we computed and stored three key parameters: the mean (*µ_cal_*), standard deviation (*σ_cal_*), and an empirical cumulative distribution function (ECDF*_cal_*).

#### Application: Real-time label generation

In the application phase, these calibrated parameters were used to generate a quantile-normalized label for each new trial. For a given trial’s raw amplitude, *a_raw_*, we first obtained its detrended value, *a_detrend_*, from the EWMA filter. This value was then standardized using the stored calibration statistics to produce a z-score, *z* = (*a_detrend_* −*µ_cal_*)*/σ_cal_*. Finally, the z-score was transformed via the stored ECDF*_cal_* to yield the final label *y* = ECDF*_cal_* (*z*). The resulting label *y* ∈ [0, 1] represents the trial’s excitability as a percentile rank relative to the initial, stable calibration distribution.

### Model architecture

The PRIME architecture (Table 1) integrates a convolutional front-end for spatiotemporal feature extraction with a state-space core for modeling long-range dependencies. The model consists of three primary components:

1. **Spatiotemporal convolutional front-end**: Inspired by DeepConvNet [27], this module uses a temporal convolution to extract initial time-domain features, followed by a spatial convolution to learn channel correlations. Pooling and dropout are applied to regularize the features and reduce dimensionality.
2. **State-space modeling**: The extracted feature sequence is processed by a bidirectional S4 block [26, 65] to capture long-range temporal dependencies within the EEG signals.
3. **Prediction head**: A final classifier, consisting of an adaptive pooling layer and a linear layer, maps the processed temporal features to a single continuous output, which is optimized using a binary cross-entropy (BCE) loss.

**Table 1.** PRIME architecture summary. The input shape corresponds to (channels, timepoints).

| Component | Output shape | Parameters |
| --- | --- | --- |
| Input | (60, 50) | — |
| Spatiotemporal front-end | (25, 25) | 37,800 |
| S4 Block | (25, 25) | 8,725 |
| Prediction head | (1) | 26 |
| **Total** |  | **46,551** |

### Training and adaptation framework

PRIME employs a hierarchical training strategy designed to achieve robust cross-subject generalization while enabling rapid online adaptation to individual neural characteristics. The framework operates through three sequential phases: population-level pretraining establishes general EEG-to-TEP mappings across subjects, subject-specific calibration adapts the model to individual neural patterns, and continuous trial-by-trial adaptation refines predictions during online deployment (see Table 2 for key hyperparameters for each phase).

**Table 2.** Training configuration across the three learning phases.

| Parameter | Population pretraining | Subject calibration | Online adaptation |
| --- | --- | --- | --- |
| **Data settings** |  |  |  |
| Subject split | 2-fold CV (50% train) | From test split | From test split |
| Trials used | All from training subjects | First 100 trials | Sliding window (50 trials) |
| **Training hyperparameters** |  |  |  |
| Epochs | 100 | 50 | 1 (per trial) |
| Batch size | 64 | 50 | 50 (window size) |
| Learning rate | $3 \times 10^{-4}$ | $1 \times 10^{-4}$ | $1 \times 10^{-4}$ |
| Warm-up trials | N/A | N/A | 50 |
| **Optimization & Loss** |  |  |  |
| Optimizer | AdamW | AdamW | AdamW |
| Loss Function | BCE with Logits | BCE with Logits | BCE with Logits |

**Table 3.**
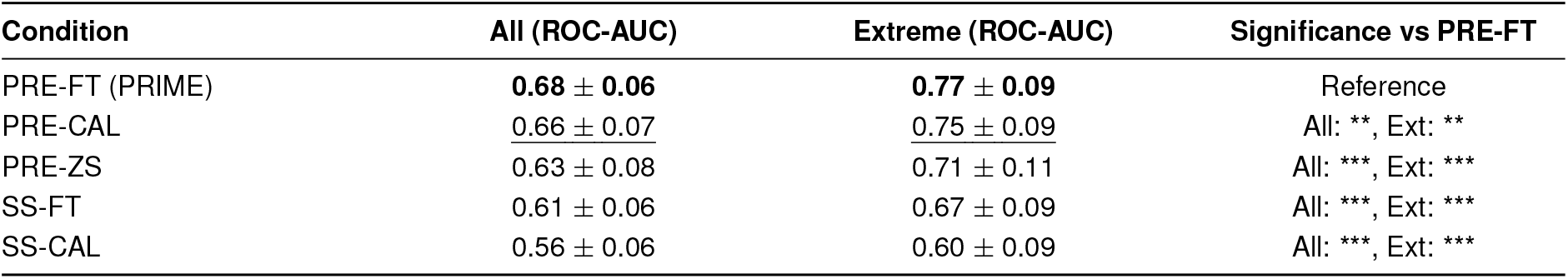
Predictive performance (ROC-AUC) across training and calibration conditions, aggregated over 50 datasets (approx. 950 trials each). Conditions include: subject-specific calibration (**SS-CAL**), subject-specific finetuning (**SS-FT**), pretrained zero-shot (**PRE-ZS**), pretrained calibrated (**PRE-CAL**), and pretrained finetuned (**PRE-FT**). Performance is shown for all trials (“All”) and for trials with the most extreme (highest and lowest 25%) TEP amplitudes (“Extreme”). Significance tests assess if PRE-FT (reference) is superior to other conditions (paired one-sided t-tests, Benjamini-Hochberg corrected).

| Condition | All (ROC-AUC) | Extreme (ROC-AUC) | Significance vs PRE-FT |
| --- | --- | --- | --- |
| PRE-FT (PRIME) | **0.68 ± 0.06** | **0.77 ± 0.09** | Reference |
| PRE-CAL | <u>0.66 ± 0.07</u> | <u>0.75 ± 0.09</u> | All: **, Ext: ** |
| PRE-ZS | 0.63 ± 0.08 | 0.71 ± 0.11 | All: ***, Ext: *** |
| SS-FT | 0.61 ± 0.06 | 0.67 ± 0.09 | All: ***, Ext: *** |
| SS-CAL | 0.56 ± 0.06 | 0.60 ± 0.09 | All: ***, Ext: *** |

**Table 4.** Comparison of PRIME against established EEG decoding models (EEGNet, ATCNet, Shallow ConvNet, Deep ConvNet) and ablated variants (PRIME w/o S4, PRIME w/ Conv). Performance is aggregated over 50 datasets (approx. 955 trials each) for all trials (“All”) and extreme amplitude trials (highest and lowest 25%, “Extreme”). Significance tests assess if PRIME (reference) is superior to other models (paired one-sided t-tests, Benjamini-Hochberg corrected).

| Model | All (ROC-AUC) | Extreme (ROC-AUC) | Significance vs PRIME |
| --- | --- | --- | --- |
| PRIME | **0.68 ± 0.06** | **0.77 ± 0.08** | Reference |
| PRIME w/ Conv | <u>0.66 ± 0.06</u> | <u>0.75 ± 0.09</u> | All: ***, Ext: *** |
| Shallow ConvNet | 0.65 ± 0.06 | 0.74 ± 0.08 | All: ***, Ext: *** |
| Deep ConvNet | 0.65 ± 0.06 | 0.73 ± 0.09 | All: ***, Ext: *** |
| PRIME w/o S4 | 0.62 ± 0.06 | 0.70 ± 0.09 | All: ***, Ext: *** |
| EEGNet | 0.60 ± 0.06 | 0.66 ± 0.09 | All: ***, Ext: *** |
| ATCNet | 0.56 ± 0.05 | 0.60 ± 0.09 | All: ***, Ext: *** |

**Table 5.** Performance comparison of different training configurations: Continual Finetuning (CFT), Euclidean Alignment + CFT (EA+CFT), Decision-level CFT (Dec-CFT), EA + Dec-CFT, EA only, and thresholded variants (Thr-CFT, EA+Thr-CFT). Significance tests assess if the standard PRIME configuration, CFT (reference), is superior to other variants across all trials and on extreme amplitude trials.

| Config | All (ROC-AUC) | Extreme (ROC-AUC) | Significance vs CFT |
| --- | --- | --- | --- |
| CFT (PRIME) | **0.68 ± 0.06** | **0.77 ± 0.09** | Reference |
| EA+ CFT | **0.68 ± 0.06** | **0.77 ± 0.08** | All: *, Ext: |
| Dec-CFT | 0.67 ± 0.07 | 0.76 ± 0.09 | All: *, Ext: * |
| EA+ Dec-CFT | 0.67 ± 0.07 | 0.76 ± 0.09 | All: *, Ext: * |
| EA | 0.65 ± 0.07 | 0.74 ± 0.09 | All: ***, Ext: *** |
| EA+ Thr-CFT | 0.63 ± 0.07 | 0.72 ± 0.11 | All: ***, Ext: *** |
| Thr-CFT | 0.63 ± 0.08 | 0.71 ± 0.11 | All: ***, Ext: *** |

**Table 6.** Impact of EEG input window size (50-500 ms) on the prediction performance of PRIME. Significance tests assess if the 50 ms window (reference) is superior to longer windows across all trials and on extreme amplitude trials.

| Window Size | All (ROC-AUC) | Extreme (ROC-AUC) | Significance vs 50 ms |
| --- | --- | --- | --- |
| 50 ms (PRIME) | **0.68 ± 0.06** | **0.77 ± 0.09** | Reference |
| 100 ms | <u>0.67 ± 0.06</u> | <u>0.77 ± 0.09</u> | All: ***, Ext: * |
| 200 ms | 0.66 ± 0.06 | 0.75 ± 0.08 | All: ***, Ext: *** |
| 300 ms | 0.65 ± 0.05 | 0.73 ± 0.08 | All: ***, Ext: *** |
| 400 ms | 0.64 ± 0.06 | 0.73 ± 0.08 | All: ***, Ext: *** |
| 500 ms | 0.64 ± 0.05 | 0.72 ± 0.08 | All: ***, Ext: *** |

**Table 7.** Performance comparison across different window locations prior to the TMS pulse (10-50 ms). Significance tests assess if the 10 ms pre-stimulus location (reference) provides superior ROC-AUC scores compared to later windows across all trials and on extreme amplitude trials.

| Window Location | All (ROC-AUC) | Extreme (ROC-AUC) | Significance vs 10 ms |
| --- | --- | --- | --- |
| 10 ms (PRIME) | **0.68 ± 0.06** | **0.77 ± 0.09** | Reference |
| 20 ms | <u>0.67 ± 0.06</u> | <u>0.76 ± 0.09</u> | All: ***, Ext: *** |
| 30 ms | <u>0.63 ± 0.06</u> | 0.70 ± 0.09 | All: ***, Ext: *** |
| 40 ms | 0.61 ± 0.06 | 0.68 ± 0.10 | All: ***, Ext: *** |
| 50 ms | 0.62 ± 0.06 | 0.69 ± 0.10 | All: ***, Ext: *** |

**Table 8.** Effect of subject-specific calibration dataset size on performance, comparing datasets of 100, 200, 300, and 400 trials. Significance tests assess if using more trials provides a performance gain over the 100-trial reference across all trials and on extreme amplitude trials.

| Calibration Trials | All (ROC-AUC) | Extreme (ROC-AUC) | Significance vs 100 Trials |
| --- | --- | --- | --- |
| 100 Trials | <u>0.68 ± 0.06</u> | 0.77 ± 0.09 | Reference |
| 200 Trials | **0.68 ± 0.06** | **0.78 ± 0.08** | All: **, Ext: *** |
| 300 Trials | **0.68 ± 0.06** | **0.78 ± 0.09** | All: **, Ext: *** |
| 400 Trials | **0.68 ± 0.07** | **0.78 ± 0.09** | All: **, Ext: *** |

**Table 9.** Performance comparison for PRIME across different physiological targets. The model’s ability to predict N45 TEP amplitudes (PRIME (TEPfree)) is the reference. This is compared against its performance when predicting a fixed orientation TEP component (TEP fixed), the P60 TEP component, and the motor evoked potential (MEP) amplitude. Significance tests assess if predicting the N45 component is superior to predicting other physiological signals across all trials and on extreme amplitude trials.

| Paradigm | All (ROC-AUC) | Extreme (ROC-AUC) | Significance vs PRIME (TEPfree) |
| --- | --- | --- | --- |
| PRIME (TEPfree) | 0.68 ± 0.06 | 0.77 ± 0.09 | Reference |
| TEP fixed | 0.68 ± 0.06 | 0.77 ± 0.08 | All: , Ext: |
| P60 (TEP) | 0.64 ± 0.06 | 0.72 ± 0.08 | All: ***, Ext: *** |
| MEP | 0.52 ± 0.04 | 0.53 ± 0.07 | All: ***, Ext: *** |

#### Phase 1: Population-level pretraining

The initial phase establishes robust EEG-to-TEP mappings using data from a large population of subjects. To enhance statistical homogeneity across subjects, we apply an optional two-step alignment process to each training subject’s data. The process first whitens each subject’s data based on its own covariance matrix, then rotates the whitened data from all subjects to a common space defined by the global average covariance of the entire training population. We employ a 2-fold cross-subject validation scheme where the model trains on half of the subjects and evaluates on the held-out half for each fold. The model undergoes pretraining for 100 epochs using the AdamW optimizer with a binary cross-entropy with logits loss function.

#### Phase 2: Subject-specific calibration

Before online deployment, the pretrained model undergoes brief calibration using an initial set of data from the target subject. The model finetunes on the first 100 trials for 50 epochs, serving two critical purposes. Model specialization allows the weights, initialized from population-level pretraining, to adapt to the specific neural patterns of the new subject. During this phase, model parameters are updated regardless of the online adaptation strategy selected for the subsequent phase. Alignment initialization uses the same 100-trial calibration set to compute an initial reference covariance matrix that seeds the optional unsupervised alignment process in the online phase, providing a subject-specific statistical baseline.

#### Phase 3: Trial-by-trial online adaptation

Following calibration, the model transitions to continuous online adaptation through three sequential steps for each incoming trial: prediction, optional unsupervised alignment, and supervised finetuning.

##### 1. Prediction

The model first generates a prediction for the current, unseen trial based on its pre-stimulus EEG features.

##### 2. Unsupervised euclidean alignment (Optional)

To investigate the utility of mitigating statistical distribution shifts between subjects, we implement an optional unsupervised alignment step. This preprocessing approach standardizes the second-order statistics of input data by whitening each trial **X** ∈ ℝ*^C×T^*(where *C* and *T* represent the number of channels and time points, respectively) as

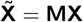

where 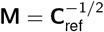 represents the inverse square root of a reference covariance matrix **C**_ref_. This reference covariance updates dynamically after each trial using an Exponential Moving Average (EMA) over the covariances of the 50 most recent trials

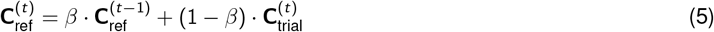

where 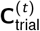 represents the covariance of the current trial and the smoothing factor *β* is set to 0.99. This alignment update occurs after prediction using only the pre-stimulus trial’s EEG data, preventing label information leakage into the adaptation process.

##### 3. Supervised continual finetuning (CFT)

After the true label for a trial is revealed, the model parameters undergo supervised updating. This finetuning begins only after collecting an initial buffer of 50 trials to ensure stable gradient estimation. Subsequently, for each new trial, the model trains for a single epoch on data within this 50-trial sliding window. We evaluate three distinct CFT strategies to understand the trade-offs between adaptability and representational stability. Full finetuning updates all model parameters, allowing the entire network to adapt to the incoming data stream and offering maximum flexibility. Decision-only finetuning preserves the learned feature hierarchy from pretraining and mitigates catastrophic forgetting by freezing feature extraction layers and updating only the parameters of the final learnable layer (either linear or convolutional) in the network. Threshold-only finetuning represents the most restrictive and computationally efficient strategy, learning only a single scalar parameter that acts as an adaptive bias applied to the model’s output logits, effectively shifting the decision threshold without altering feature extractor or classifier weights.

#### Evaluation metrics

We assess model performance using the Area Under the Receiver Operating Characteristic Curve (ROC-AUC) due to its robustness to class imbalance. To dissect the contributions of each framework component, we evaluate performance at three distinct stages that isolate different aspects of the learning process.

Pretrained zero-shot (PRE-ZS) provides a baseline for out-of-the-box generalization by applying the general population model directly to all trials from a new subject without any subject-specific calibration or finetuning. Pretrained calibrated (PRE-CAL) isolates the benefit of the initial calibration phase by evaluating model performance on remaining trials after calibration on the first 100 trials, without further online adaptation. Pretrained finetuned (PRE-FT) represents the final performance of the complete framework, measuring overall ROC-AUC across all predictions made during continuous, trial-by-trial online adaptation following calibration.

For each stage, we report performance on two data subsets to provide comprehensive evaluation: across all available test trials (“All”) and specifically for trials with extreme TEP amplitudes (highest and lowest 25% of responses). Extreme trials represent the most clinically relevant scenarios where clear distinctions between high and low excitability states are crucial for therapeutic applications. The final reported scores for each condition are averaged across all subjects and test folds.

### Experiments

Our experimental evaluation was designed to: 1) systematically assess the performance of the full PRIME framework against ablated and alternative conditions; 2) benchmark it against established deep learning architectures; 3) analyze its sensitivity to key hyperparameters; and 4) test the generalizability of its learned representations to different physiological targets. To determine if our proposed framework significantly outperformed the alternatives, we compared the performance of PRIME against each other condition using a paired, one-sided t-test. P-values were corrected for multiple comparisons across the datasets using the Benjamini-Hochberg procedure (* p < 0.05, ** p < 0.01, *** p < 0.001)

#### Ablation and comparison conditions

To systematically evaluate the contribution of each framework stage, we compared the full PRIME system against a hierarchy of ablated and alternative conditions. Each condition isolates the impact of population-level pretraining, subject-specific calibration, and continual finetuning through controlled comparisons.

**Subject-specific calibration only (SS-CAL)** establishes a baseline for performance using only a small, subject-specific dataset. The model initializes with random weights without pretraining and trains exclusively on the 100-trial calibration set. Model weights are then frozen and performance is evaluated on remaining trials. **Subject-specific finetuning (SS-FT)** represents a “from-scratch” learning approach that omits population-level pretraining. The model initializes with random weights and follows the same two-phase training protocol as the full framework: initial training on the 100-trial calibration set followed by trial-by-trial continual finetuning on all remaining data.

**Pretrained zero-shot (PRE-ZS)** serves as the primary baseline for generalization from the population model by applying the pretrained model directly to new subject data without calibration or finetuning, with weights remaining frozen during evaluation.**Pretrained + calibrated (PRE-CAL)** isolates the benefit of the calibration step by updating the population-pretrained model using the 100-trial subject-specific calibration set, then freezing weights and evaluating performance on remaining trials without continual finetuning.

**Pretrained + finetuned (PRE-FT)** represents the complete proposed PRIME framework, leveraging all three stages: model initialization from population-level pretraining, specialization during subject-specific calibration, and continuous updating through trial-by-trial finetuning on all subsequent data.

#### Benchmark architectures

To contextualize the performance of PRIME, we benchmarked it against four established deep neural network architectures from the EEG decoding literature. Given the short input sequences used in our task (50 time points, corresponding to 50 ms of data), we appropriately scaled down the kernel sizes and pooling layers of the original benchmark architectures to prevent excessive temporal dimension reduction and ensure fair comparison. **ShallowConvNet** provides a compact architecture inspired by the filter bank common spatial pattern (FBCSP) pipeline [27], employing temporal convolution followed by spatial filtering to extract spatiotemporal features. **DeepConvNet** implements a deeper, multi-layer architecture composed of stacked convolutional blocks [27], designed to learn feature hierarchies from low-level signal characteristics to more abstract and discriminative representations.**EEGNet** offers a highly compact and efficient architecture that utilizes depthwise and separable convolutions to learn temporal and spatial filters independently [**Lawhern2018**]. **ATCNet** presents a hybrid architecture that augments a convolutional frontend with multi-head self-attention and a temporal convolutional network (TCN) [29], enabling the model to capture long-range temporal dependencies and explicitly weight the importance of learned features.

#### Architectural ablations

To isolate the contribution of the Structured State Space (S4) component within the PRIME architecture, we designed and evaluated two ablated variants that systematically remove or replace this core component.

**PRIME without S4** removes the entire S4 temporal modeling block, leaving the model to consist solely of the spatiotemporal convolutional frontend feeding directly into global average pooling and the final linear classifier. This ablation measures performance attributable to the convolutional feature extractor alone. **PRIME with convolutional core** replaces the S4 block with a standard temporal feature extractor comprising a stack of conventional 1D convolutional layers of equivalent depth, enabling direct comparison between S4’s ability to model long-range dependencies and traditional CNN-based approaches.

#### Hyperparameter sensitivity analysis

We investigated the sensitivity of PRIME to three key hyperparameters using the full PRE-FT configuration as baseline. We varied the number of trials used for subject-specific calibration, testing the default 100 trials against 200, 300 and 400 trials to assess calibration requirements. We evaluated the impact of pre-stimulus EEG window size by testing windows of 50 ms (default), 100 ms, 200 ms, 300 ms, 400 ms, and 500 ms to determine optimal temporal resolution. We examined the importance of window temporal location by testing offsets ending 10 ms (default), 20 ms, 30 ms, 40 ms and 50 ms before the TMS pulse to identify the most predictive temporal window placement.

#### Generalizability to different physiological targets

To assess whether features learned by PRIME generalize beyond predicting the N45 component, we conducted two additional experiments using the best-performing PRE-FT configuration. We tested intra-domain generalizability by retraining the framework to predict the later P60 TEP component, examining whether similar cortical excitability mechanisms underlie different TEP responses. We evaluated cross-domain generalizability by repurposing the framework to predict MEPs as a measure of corticospinal excitability derived from EMG signals, to determine whether EEG features predictive of cortical responsiveness extend to downstream motor pathway excitability.

### Real-time latency analysis

To assess the real-time feasibility of PRIME, we quantified the computational latency of its core online components under realistic hardware constraints. All benchmarks were conducted in a PyTorch environment [66] using an 8-core CPU (AMD EPYC 7302) and a consumer-grade GPU (NVIDIA GeForce RTX 2080Ti with 11 GB VRAM) to reflect standard laboratory computing setups.

We measured wall-clock time in milliseconds for four distinct computational stages across all subjects, grouping operations by their occurrence relative to the TMS pulse. For GPU benchmarks, we ensured accurate timings by synchronizing the device before and after each measurement to account for asynchronous operation completion. Prediction and model adaptation measurements were averaged over 100 repetitions, preceded by 10 warm-up runs to stabilize system state and eliminate initialization overhead.

#### Pre-stimulus operations

These operations must complete within the interval between EEG recording and TMS pulse delivery to maintain real-time performance. EEG preprocessing encompasses the time required to process incoming pre-stimulus EEG segments, including filtering, average-referencing, and threshold-based trial rejection. TEP prediction measures the time for a single forward pass of the PRIME model to generate predictions from preprocessed EEG data.

#### Post-stimulus operations

These operations occur during the inter-stimulus interval after TMS pulse delivery, allowing more relaxed timing constraints. Post-stimulus processing and TEP extraction quantifies the time required for all post-pulse data handling, encompassing initial processing of post-stimulus EEG data and subsequent extraction of the relevant TEP component (e.g., N45) that serves as the prediction target for model adaptation. Model adaptation measures the time to perform one complete supervised finetuning step, including forward pass, loss calculation, backpropagation, and optimizer weight update using a batch of the 50 most recent trials, consistent with the online learning protocol.

#### Neurophysiological feature interpretability

To identify the neurophysiological features underlying the predictive capabilities of PRIME, we conducted occlusion-based sensitivity analysis on final, subject-specific finetuned models. This approach systematically removes distinct spatial or spectral components from input EEG signals and quantifies the resulting degradation in model performance, measured as the change in ROC-AUC (Δ ROC-AUC). Larger performance drops indicate higher importance of the occluded feature. We assessed statistical significance of performance drops using one-sided Wilcoxon signed-rank tests, with p-values corrected for multiple comparisons using the Benjamini-Hochberg procedure. Three complementary analyses created a comprehensive interpretability map of the decision-making process of PRIME.

##### Spatial importance mapping

To identify spatial origins of predictive information, we occluded broadband EEG signals from each electrode and its eight nearest neighbors. The resulting performance drop for each occlusion was mapped to the corresponding electrode location, creating topographical maps of spatial importance across the scalp. This analysis reveals which cortical regions contribute most critically to excitability predictions.

##### Frequency band importance

To determine the contribution of distinct neural oscillations, we selectively removed individual frequency bands across all electrodes using band-stop filters (4th order Butterworth). We evaluated four canonical bands: theta (4–8 Hz), alpha (8–13 Hz), beta (13–25 Hz), and gamma (25–47 Hz). The aggregate drop in ROC-AUC for each occluded band reflects its overall predictive value for cortical excitability assessment.

##### Spatially resolved frequency analysis

To create detailed spatio-spectral profiles, we combined spatial and frequency approaches by systematically occluding each frequency band at specific electrode locations (including eight nearest neighbors). This procedure mapped the precise cortical locations from which predictive oscillatory activity originates, providing spatially and spectrally resolved feature importance maps that reveal the neurophysiological basis of the excitability predictions of PRIME.

## Data availability

De-identified data is available from the corresponding authors upon reasonable request, subject to ethical approval and data agreements.

## Code availability

Analysis code along with relevant documentation is publicly available on GitHub at https://github.com/mackelab/PRIME/

## Ethics declarations

The authors declare no competing financial or non-financial interests.

## Supplementary results

**Figure S1.**
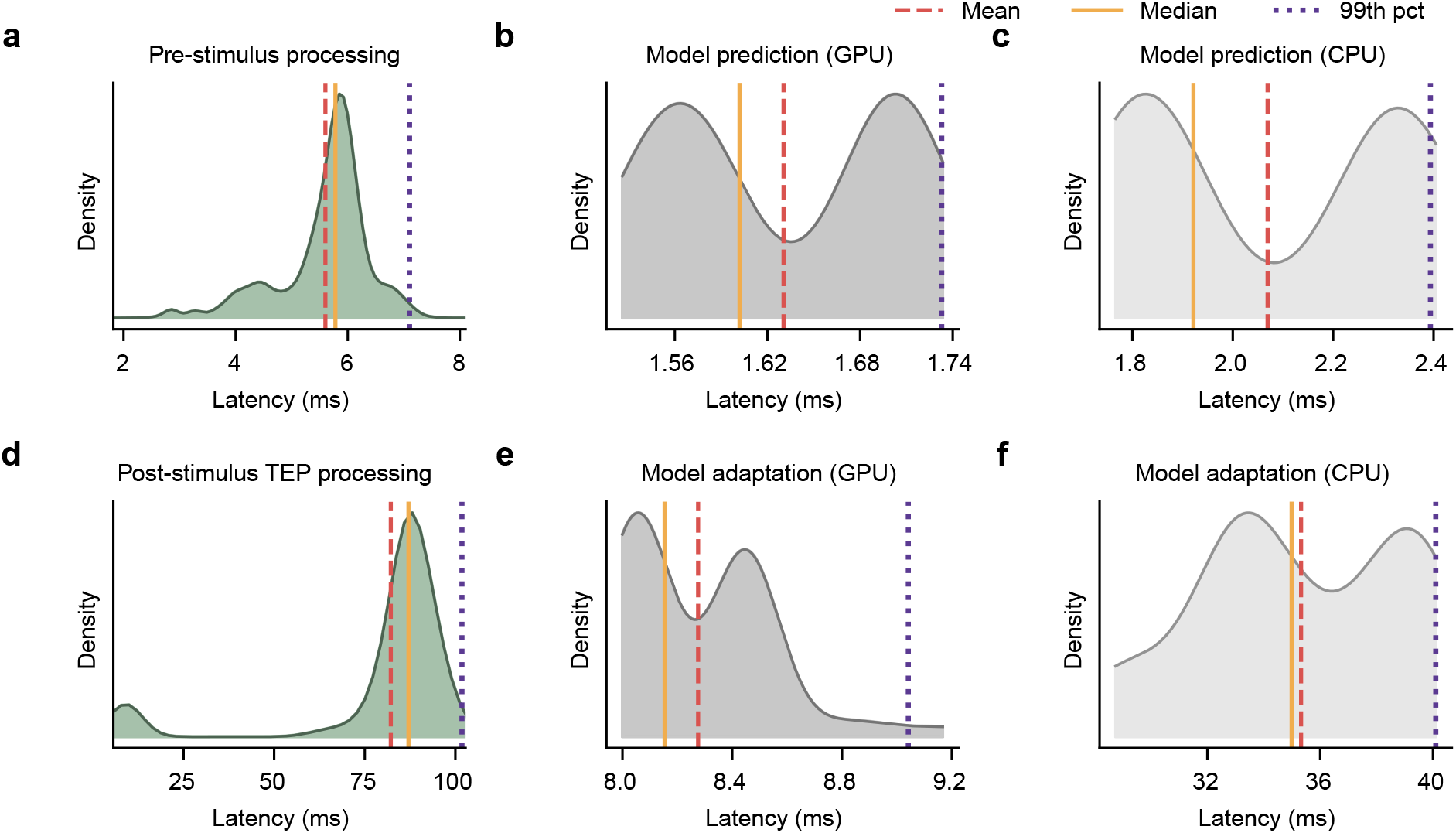
Computational latency analysis for real-time PRIME framework implementation. Smoothed histogram plots (Kernel Density Estimates) illustrate the distribution of processing times for key computational stages on both an 8-core CPU and NVIDIA 2080Ti GPU. Each panel displays the latency distribution, with vertical lines marking the mean, median, and 99th percentile. **a.** Latency for CPU-based pre-stimulus EEG preprocessing, encompassing all signal cleaning and artifact rejection steps prior to the TMS pulse. **b, c.** Latency for TEP amplitude prediction using the pretrained PRIME model on GPU (**b**) and CPU (**c**). **d.** Latency for CPU-based post-stimulus processing, which includes TEP feature extraction and dipole source fitting. **e, f.** Latency for a single-epoch model adaptation (finetuning) step on GPU (**e**) and CPU (**f**).

**Figure S2.**
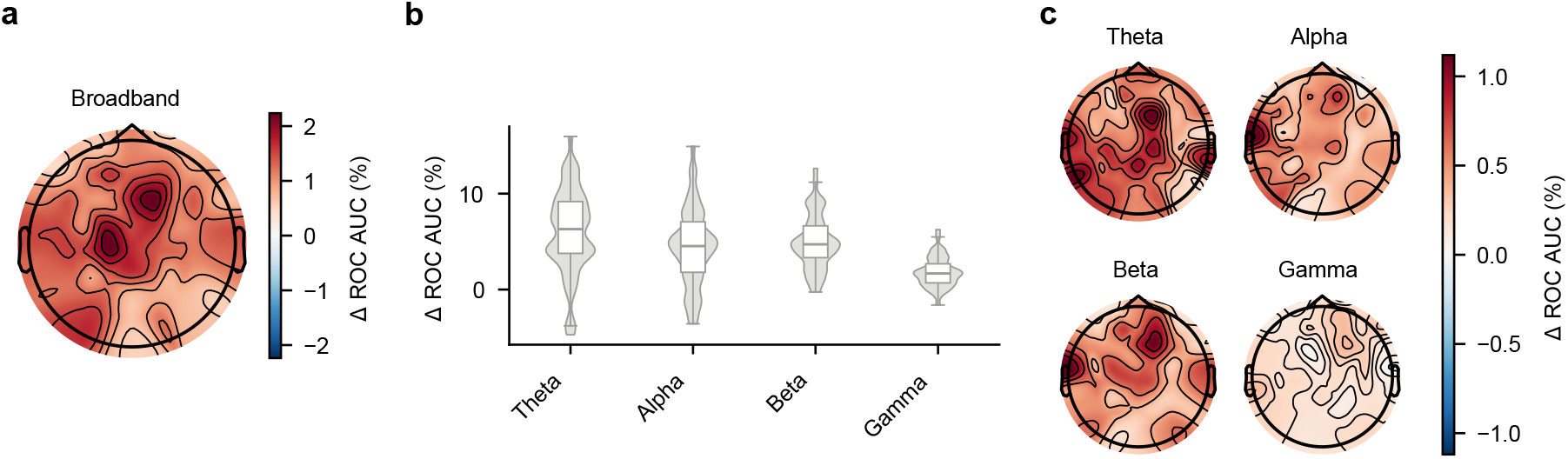
Theta and alpha oscillations in fronto-central regions provide the primary predictive features for the P60 component of TEPs. **a.** Spatial distribution of electrode importance determined by occlusion analysis. Prediction accuracy decreases (positive Δ ROC-AUC values, %) show the importance of individual electrodes and their 8-nearest-neighbors for broadband EEG-based prediction. **b.** Frequency-band importance analysis shows the contribution of theta, alpha, beta, and gamma bands across all electrodes. Positive Δ ROC-AUC (%) indicates the importance of each frequency band for accurate prediction. **c.** Spatially resolved frequency-band contributions reveal localized predictive patterns obtained by selectively occluding frequency bands at individual electrodes (8-nearest-neighbor approach). Positive Δ ROC-AUC (%) indicates the importance of specific frequency bands at different cortical locations. All analyses used subject-specific finetuned PRIME models, with statistical significance determined using Wilcoxon signed-rank tests with FDR correction across test subjects (all displayed results significant at p < 0.05).

## Notes

https://github.com/mackelab/PRIME

### Competing Interest Statement

The authors have declared no competing interest.

